# SEQSIM – A novel bioinformatics tool for comparisons of upstream gene regions – a case study of calcium binding protein spermatid associated 1 (CABS1)

**DOI:** 10.1101/2024.05.03.592313

**Authors:** Joy Ramielle L. Santos, Weijie Sun, A. Dean Befus, Marcelo Marcet-Palacios

## Abstract

The regulation of gene expression is carefully overseen by upstream gene regions (UGRs) which include promoters, enhancers, and other regulatory elements. Understanding these regions is difficult using standard bioinformatic approaches due to the scale of the human genome. Here we present SEQSIM, a novel bioinformatics tool based on a modified Needleman-Wunsch algorithm that allows for fast, comprehensive, and accurate comparison of UGRs across the human genome.

In this study, we detailed the applicability and validity of SEQSIM through an extensive case study of the calcium binding protein spermatid-associated 1 (CABS1). By analyzing 2000 base pairs upstream of every human gene, SEQSIM identified distinct clusters of UGRs, revealing conserved motifs and suggesting potential regulatory interactions. Our analysis identified 41 clusters, the second largest of which contains the CABS1 UGR. Studying the other members of the CABS1 cluster could offer new insights into its regulatory mechanisms and suggest broader implications for genes involved in similar pathways or functions.

The development and implementation of SEQSIM represents a significant step forward for the genomics field, providing a powerful new tool to dissect the complexity of the human genome and gain a better understanding of how gene expression is regulated. The study not only shows that SEQSIM is an effective means to identify potential regulatory elements and gene clusters, but also opens up new lines of inquiry to understand overall genomic architecture.

## 1. Introduction

The regulation of gene expression is an essential and complex process. The regulatory nucleotide sequences adjacent to genes include promoters, enhancers, silencers, and other elements that determine when a gene is expressed. Promoter elements are sections of the upstream gene regions (UGRs) that interact with RNA polymerases and other transcription factors to initiate transcription. Enhancers, on the other hand, are more distant regulatory elements that can activate or enhance the action of promoters. Silencers are regions that can inhibit gene expression by blocking the interaction between transcription factors and promoters. Other regulatory elements such as insulators, boundary elements, and non-coding RNAs add further complexity to the regulation of gene expression(1,2). Although all human genes are differentially regulated, there are common regulatory elements used according to gene type e.g. inducible vs. house-keeping genes. By studying and grouping these regulatory elements together, we could learn more about the genetic mechanisms that drive different functional outcomes.

In an effort to understand human promoter sequence diversity, Gagniuc et al. performed sequence homology analysis of 8512 promoter sequences available in a promoter sequence database(3). The authors categorized promoters into 10 groups according to a 2-dimensional analysis consisting of plots of Kappa Index of Coincidence vs C+G % within the -499 to +100 positions relative to the transcription initiation site. This paper suggested categorizing the promoters according to the presence of promoter elements (TATA box, GC-box and CCAAT-box), DNA curvature, or nucleosome positioning(4–9). Unfortunately, this approach is limited by the simplistic nature of 2D comparisons, leading to strictly tabular analysis of promoter types, and is unable to group promoters based on predicted functional families. Another limitation of this study is that they analyzed a small number of promoters available in databases at the time. Additionally, we know that there are important functional elements further upstream e.g. (-1950 to -522)(10). To overcome these limitations, we performed a comprehensive homology analysis of 2000 base pairs upstream of every human gene, thus including promoter proximal (-522) and middle (-1950) regions. To address these challenges, our method distinctively employs sequence similarity analysis using a modified Needleman-Wunsch algorithm(11). This adaptation enhances the ability of the software to identify and categorize UGRs.

We hypothesize that a deeper understanding of the regulation of gene expression can be attained by comparing the sequence homology of all UGRs in the human genome, then clustering them based on homology. For instance, selected genes involved in a particular type of immune response could be grouped based on similarities in their UGRs, which may contain conserved motifs recognized by the same set of regulatory elements. Similarly, genes active during specific developmental stages might have UGR elements that respond to the same developmental signals. UGRs within each cluster could then be further analyzed to discover conserved regulatory elements, their locations within the UGRs and their relationships. It is possible that a conserved enhancer element might be found in the UGRs of multiple genes in a cluster related to a specific metabolic process. Understanding the position and the sequence of such an enhancer could provide insights into how these metabolic genes are co-regulated.

Unfortunately, commonly used tools, like Clustal Omega, often have file size restrictions that limit full-genome dataset analysis. Indeed, we attempted to import a genome-wide dataset using Clustal Omega but was unsuccessful due to the 4MB size limitation. Parsing the dataset still resulted in analysis times that were prohibitively slow and estimated to take more than one year to complete. There is an obvious need for a fast and efficient tool that can compare UGRs across the entire genome. Accordingly, we developed and tested a new algorithm and software tool, SEQSIM (Sequence Similarity), which enables fast, pairwise comparison of the UGR of every human gene in the Genome Reference Consortium Human Build 38 (GRCh38.p14).

SEQSIM could be especially useful in the study of unannotated genes. We selected calcium binding protein spermatid-associated 1 (*CABS1),* a poorly characterized gene, as our pilot. The CABS1 protein is most abundant in the testis, and found in salivary glands(12). Recent analysis of GeoData has detected *CABS1* mRNA widely expressed in human tissues(13). To date, CABS1 has been associated with spermatogenesis in several species and with acute psychological stress in humans(14–17). A 7 amino acid peptide sequence in human CABS1 near its carboxyl terminus has anti-inflammatory activity. Previously, we conducted other *in silico* techniques to better understand CABS1 and its behavior(18). Using *CABS1* as a case study presents a unique challenge and opportunity as it does not have another homologous human gene or known protein isoforms. Thus, a SEQSIM analysis of *CABS1* UGR and identification of other genes with similar UGR sequences may help reveal other functions of CABS1 in various tissues and the mechanisms that regulate its expression.

## 2. Methods

SEQSIM analyzes the human genome in under an hour, achieving 27 million comparisons per minute. The software operates on C++ and is compatible with both Linux and Windows operating systems, importing sequences to compare against each other to produce a homology matrix. The pipeline involves: data mining of genome data using a Python script; utilization of a specialized scoring function to draw comparisons between sequences of 2000 nucleotides; homology matrix visualization and data clustering using various programs; and validation using 3^rd^ party software. This pipeline is summarized in Figure 1.

**Figure 1.**
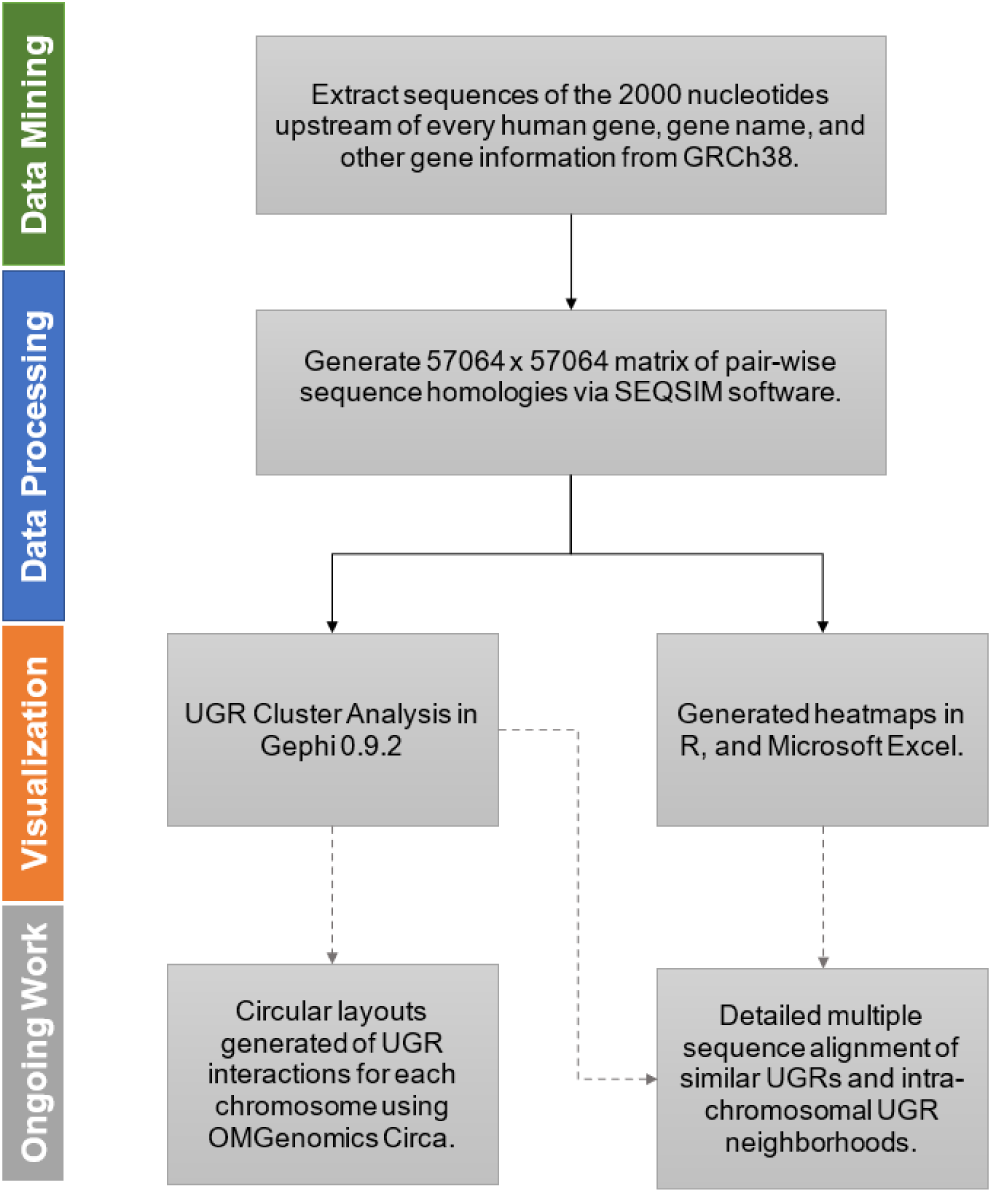
Methodology Pipeline using the novel comparison software generated for this study. The methodology occurred in three distinct phases, data mining, data processing, and data visualization approaches. Ongoing and future approaches are also denoted by the dashed arrows.

### 2.1 SEQSIM Software Development

The software, *SEQSIM* (Sequence Similarity), was created to perform DNA sequence similarity analyses at a rate of 27 x 10^6^ comparisons per minute, completing a full human genome analysis in under 1 hour. SEQSIM imports a list of named sequences and compares all sequences against each other to generate a sequence homology matrix with a percent similarity based on our comparison algorithm.

The data extraction software is written in the C++ language and can be run in a Linux or Windows operating system.

#### 2.1.1 Data Mining and Preparation of Comma Separated Values (CSV) Import File

To prepare the input files required for SEQSIM, a data mining python script was used to extract information from the GRCh38. Chromosome information was downloaded in text format from the NCBI (https://www.ncbi.nlm.nih.gov/gene) with the corresponding chromosome search criteria, e.g. `1[CH] AND human[ORGN]`. The script then extracted information such as the nominal gene symbol and name used to reference the upstream gene regions (UGRs), gene type, gene chromosomal location and coordinates, directionality, gene description and gene exon count. Next the chromosome sequence was downloaded from NCBI’s GenBank repository (19). NC_000001, refers to chromosome 1. Each chromosome FASTA sequence was downloaded separately. The first nucleotide of each gene and gene orientation was then used to extract the 2000 nucleotide UGRs in the 5’ to 3’ direction. The program compiled all this information into a commo separated values (CSV) file.

For SEQSIM, an additional CSV file was created with the identification of the sequence in the first column and the sequences to be analyzed in the next column. All sequences were required to be equal in size and were imported into the program as a string, a sequence of non-numerical characters.

#### 2.1.2 Comparison Algorithm

The program currently defines a pairwise similarity score, normalized in the range of 0 and 1 between two nucleotide sequences or strings of the same size (N = 2000 nucleotides). An anchoring sequence is identified, X, and aligned to a second sequence, Y (comparator sequence). When both sequences are perfectly aligned so that both 2000 nucleotide (i.e.: N = 2000) sequences are squared to each other, we call this position 1 (𝑖). At position 1 (𝑖 = 1), the program finds the longest string of nucleotides within the anchoring sequence that corresponds exactly to a segment in the comparator sequence. A match of 1 nucleotide corresponds to a value of 1 and each subsequent nucleotide match adds 1 point. When a mismatch is discovered, implying that the two sequences are not identical, the comparator sequence Y is shifted left by one nucleotide (ie: comparing position 1 of the anchoring sequence with position 2 of the comparator sequence) and another comparison check is conducted at this second position. It’s important to note that despite detecting mismatches, SEQSIM still compares sequences downstream of mismatched nucleotides. This process repeats until 2000 comparisons have been made between the anchor sequence X and the entire comparator sequence Y, creating a row of data that identifies the length of the segment in the comparator sequence that exactly matches a segment in the anchor sequence. This process is repeated for positions 2 to 2000 of the anchoring sequence to generate 1999 more rows of data and 4,000,000 data points for further processing (see Figure 2 and Supplemental Figure 1 for a more comprehensive example).

**Figure 2.**
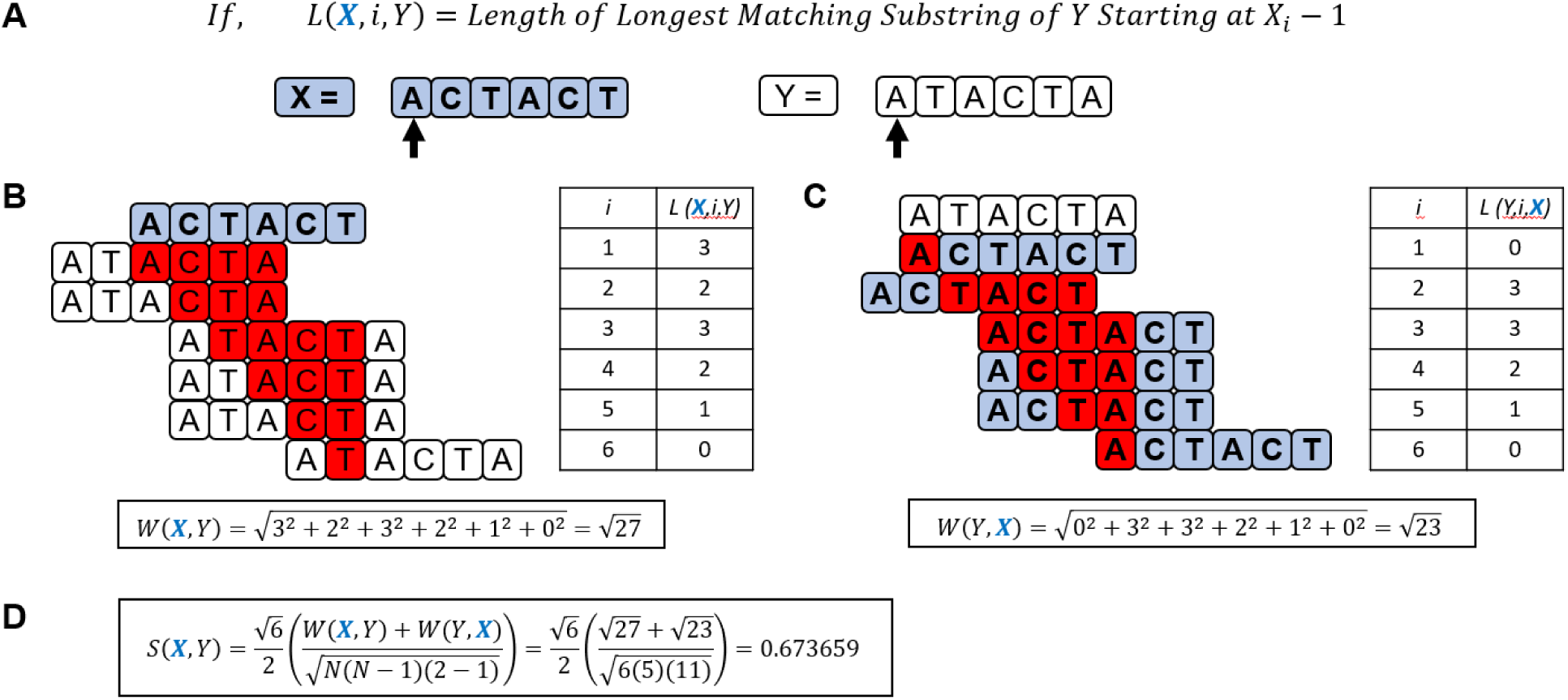
Case study conducted to develop the scoring algorithm used in our software. (A) Defines the variables and sequence notation used in our algorithm in a sample case study using the sequences X = ACTACT and Y = ATACTA. Variable L indicates the length of the longest matching substring of the comparator sequence minus one. The variable ‘i’ indicates the position on the first nucleotide of the first sequence (the reference sequence) in the notation, L(X,Y). Position 1 on Sequence X and Y is indicated by the black arrow. (B) The largest substring match, highlighted in red, in Sequence Y when compared to each position, i, in Sequence X minus 1, or L(X,i,Y). A weighing function is applied, squaring the maximum L(X,i,Y) and summing the values. (C) The largest L(Y,i,X) with the same weighing function applied. (D) The values calculated in Panels (B) and (C) substituted into the overall scoring function (S) used in our algorithm with a value between 0 to 1.

In the resulting array, the maximum value generated from each positional comparison on the anchor sequence is then carried into our scoring algorithm (See Supplementary Materials for detailed math functions). The scoring function assigns a value between 0 and 1, which can then be converted into a percentage. Higher scores are assigned to sequences that share long segments of matching nucleotides.

#### 2.1.3 User Interface

Because *SEQSIM* software is written in C++, reference files, such as the GRCh38 dataset, need to be placed in the same directory out of which the code is run. A sample of the code can be found in the Supplementary Materials.

### 2.2 Cluster Analysis

Cluster analysis of UGR similarity was performed using the open-source, network visualization software, Gephi (version 0.9.2). Data used in the analyses consisted of a set of nodes, the UGR gene ID, and edges, their pairwise similarity score. To visualize the network, Gephi’s continuous graph layout algorithm, ForceAtlas2 was implemented due to its ability to graph clusters in large datasets with high precision and speed. The algorithm used a force-directed approach to layout the nodes in 2D space, with nodes that were more similar (higher S) being drawn closer together. ForceAtlas2 also utilizes a gravity function that allows nodes to stay clustered and prevent them from drifting apart(20).

Once the data was imported to Gephi, default settings for ForceAtlas2 were used, including a gravity value of 1.0. To generate images, the “Prevent Overlap” option under Behavior Alternatives was enabled. ForceAtlas2 was run until convergence was achieved. Nodes that did not appear to cluster were filtered using modularity filters, such as degree distribution. The filter range parameters displayed nodes within the range of 11 to 502.

### 2.3 Heatmap Generation

Heatmaps were first generated using RStudio, a popular open-source statistical software (R 3.3.0+). Inputs for each heatmap consisted of a matrix of numerical values (the similarity scores, S) between UGRs within a specific chromosome in sequential order. The color scale chosen represented the range of values within the data set, red being a high similarity score and white a low score.

Subsequent heatmaps were generated using conditional formatting in Microsoft Excel. Each chromosome matrix was imported into Excel. The conditional formatting tool was then applied. Under the “highlight cell rules” option, a custom rule was created that utilized a 3-color scale. Depending on the chromosome, the color scale could be adjusted to match the specific data being analyzed. As with the heatmaps generated in R, red indicated high similarity scores and white low scores.

### 2.4 Validation with Clustal Omega and Multiple Sequence Alignment

The UGR sequences used in SEQSIM were exported into FASTA format. The resulting file was imported into the online server for Clustal Omega, a widely used tool for multiple sequence alignment (MSA). The resulting Percent Identity Matrix generated using Clustal Omega was then visualized using the methods described in Section 2.3. For some datasets, Clustal Omega automatically generated an identity matrix with a different order of UGRs. Rearrangements of the data were conducted in Microsoft Excel.

Another automatically generated output in Clustal Omega was a .CLUSTAL_NUM alignment file. This file was converted to FASTA format using the Clustal to Fasta Sequence Converter by Bugaco.com. Similar UGR matches to that of CABS1, or sequences of interest were visualized using NCBI’s Multiple Sequence Alignment Viewer 1.23.0 by uploading the resulting FASTA file to the server. CABS1 UGR, our case study gene, was then set as the “anchor” so all other sequences were compared relative to CABS1’s UGR. The other sequences were also arranged from highest to lowest homology to the anchor. For clarity, the resulting alignment was colored green for matching nucleotide sequences and red for differences. Any large regions of interest were searched in NCBI’s nucleotide BLAST (Basic Local Alignment Search Tool) server.

## 3. Results

### 3.1 SEQSIM Generated Score Matrix and Cluster Map

SEQSIM generated a 57064 x 57064 matrix of similarity scores between the UGRs of every human gene. Due to computational restraints, we only performed a Gephi cluster analysis of approximately the top 10% most similar UGRs of the complete matrix. This cluster analysis identified five main clusters with more than 100 UGRs (nodes), with the third largest containing CABS1 (See Figure 3, Panel B). Using the Louvain method to determine modularity, Gephi generated a score of 0.840 and discovered 41 total communities or clusters (See Figure 3C). The largest cluster had 402 UGR members and the smallest just 1 member.

**Figure 3.**
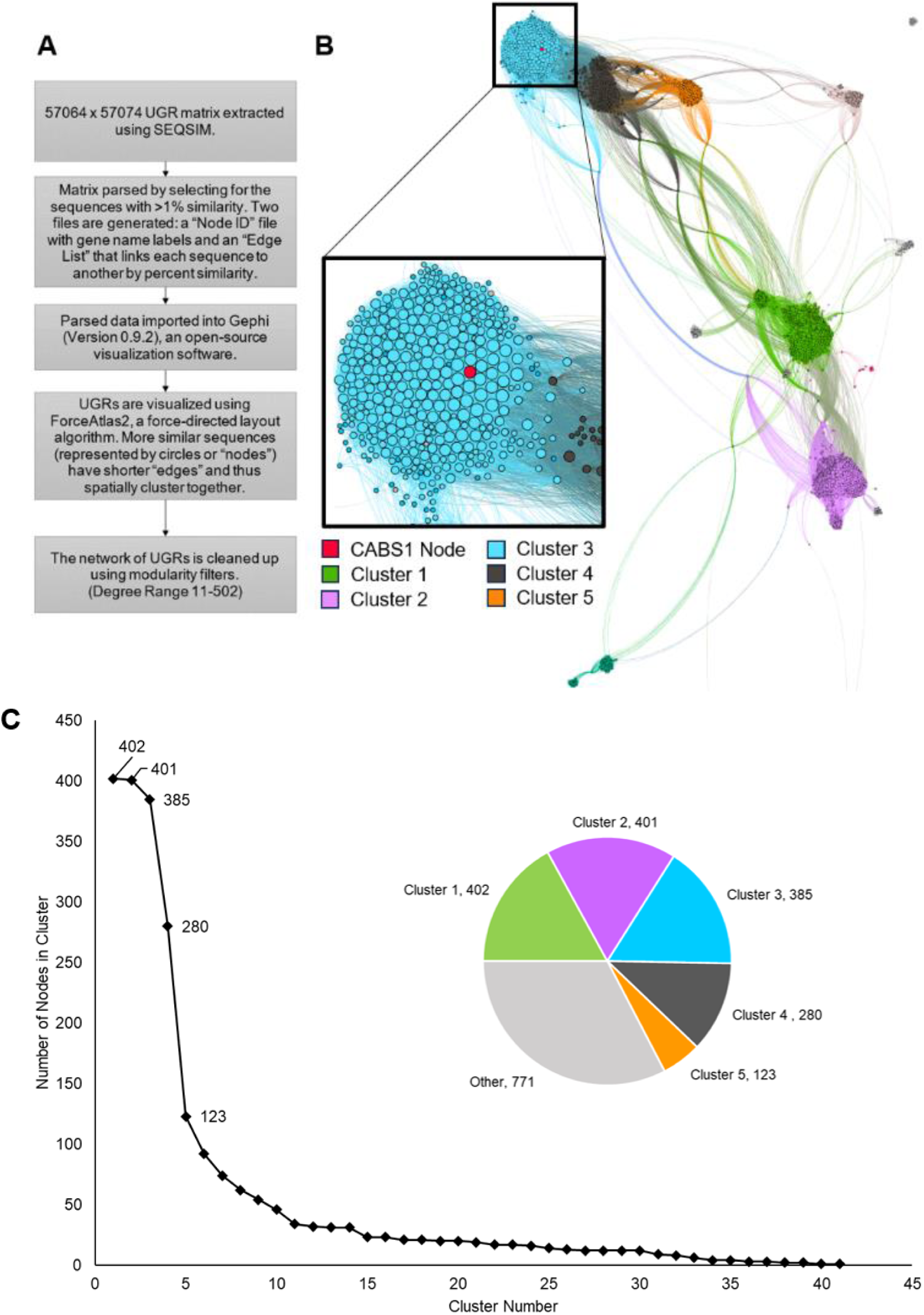
Cluster analysis of the top 10% most similar UGRs of the human genome. (A) Methodology Pipeline for generating the cluster diagrams in Gephi 0.9.2. 10% of our 57064 x 57604 large matrix was visualized using Gephi 0.9.2, an open-source data visualization software. Each upstream gene region (UGR) is depicted in the software as a circle, or a ‘node.’ When there is similarity between sequences, a line, also known as an ‘edge,’ is drawn between each sequence. The more similar the sequence, the shorter the edge. The clusters were generated using Gephi’s ForceAtlas2 algorithm. CABS1 can be seen, labelled red. (B) Depicts the network diagram of the topmost similar UGRs mined from the GRCh38. Due to computational restraints, only 10% of the analyzed UGRs can be displayed. From the analysis, 5 main clusters were discovered colored blue, green, pink, black and orange. Several other smaller, clusters were also observed. A zoomed view of cluster 3 shows the CABS1 UGR in red. Nodes are sized according to the number of other UGRs to which they are similar. The higher the number of hits, the larger the node. CABS1, with its multiple connections to many other UGRs, appears to be a large contributor to its cluster. (C) Characterization curve of clusters in the human genome. Over 41 clusters were discovered with varying populations of 1 to 402 nodes.

After filtering out UGRs with no connecting edges, 2362 nodes remained, 67.4% of which were within the main 5 clusters. The CABS1 UGR cluster comprised of 385 members, with 258 members directly connected to CABS1. These included many UGRs for non-coding RNA and pseudo genes. According to our algorithm, the UGR of CABS1 (red node in Fig 3B) is highly similar to the UGRs of the protein-coding genes VWCE, SPOCK1, THSD4, RNF39, and TMX2 with similarity scores of 5.16%, 4.89%, 4.03%, 4.01%, and 3.80% respectively. Our scoring algorithm is very strict and does not allow for gaps or substitutions that other programs, like Clustal Omega, allow. When these same UGRs are compared using Clustal Omega, scores were 78.70%, 81.99%, 82.89%, 96.54%, and 77.09% for the same gene UGRs. Interestingly, these 5 genes are found on different chromosomes than CABS1 (see Table 1).

**Table 1.**
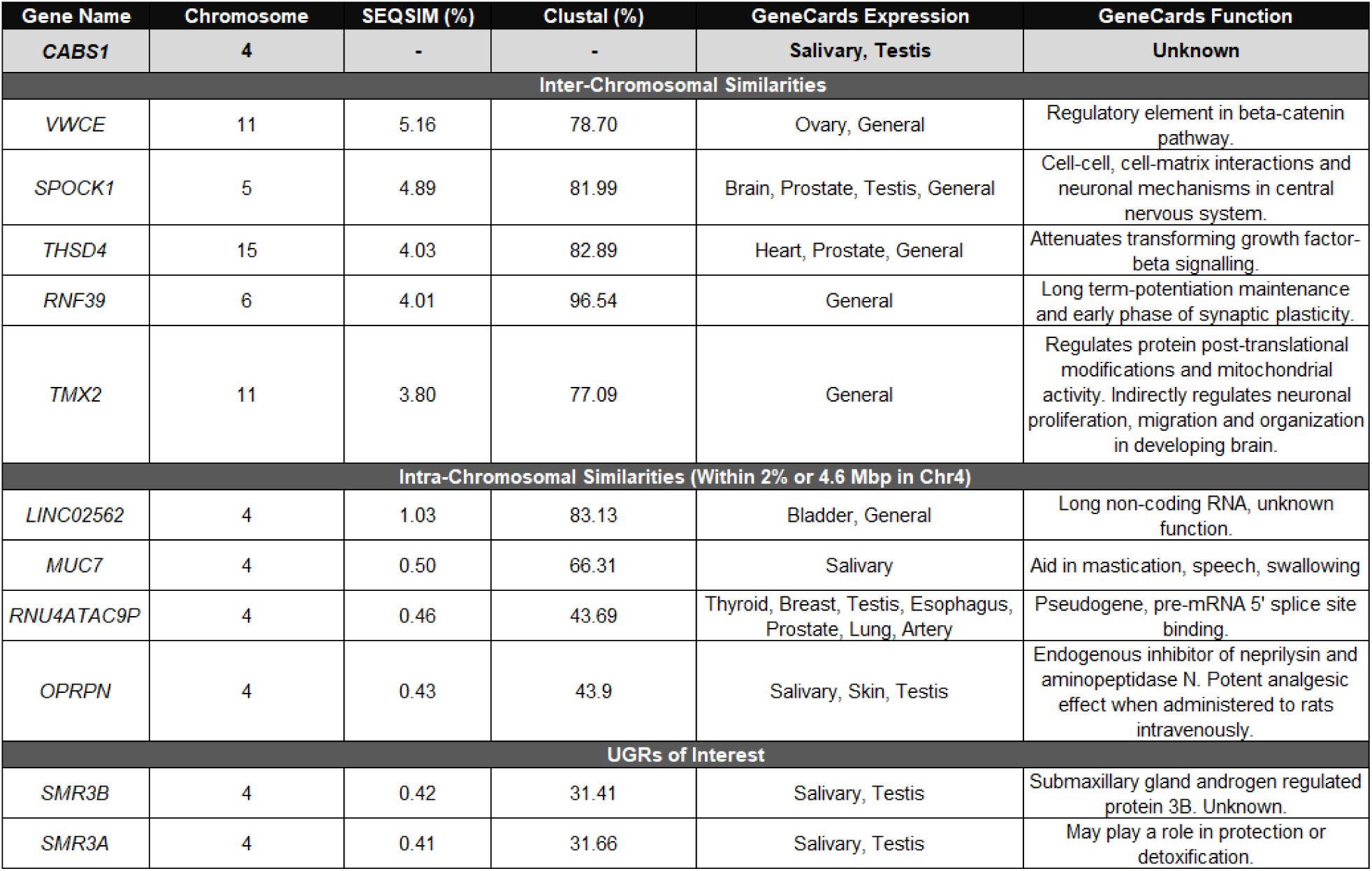
Summary Table of the UGRS of interested related to CABS1. UGRs in the Inter-Chromosomal and Intra-Chromosomal sections had high similarity UGRs to that of CABS1. Inter-Chromosomal UGRs were found to be clustered with CABS1 in our SEQSIM analysis. Also in the table, are SMR3A and 3B, which did not cluster with CABS1 and did not show high similarity in UGRs.

### 3.2 Heatmap of Adjacent Genes around Case Study UGR of CABS1 in Chromosome 4

The heatmap generated for 200 UGRs flanking the CABS1 gene loci in the same order as UGRs appear in the DNA is shown in Figure 4A. Within Chromosome 4, and within 50 genes on either side of CABS1, the UGRs for LINC02562, MUC7 and UGT2B11 closely resemble the UGR of CABS1. Other matches included genes that have not yet been defined (ie: LOC genes). Comparative validation studies in Clustal Omega (Figure 4B) resulted in new potentially similar UGRs in addition to those previously discovered using SEQSIM. This included the UGR of RNU4ATAC9P and the OPRPN protein coding gene which like CABS1 is associated with male reproductive function among other roles(21), see Table 1.

**Figure 4.**
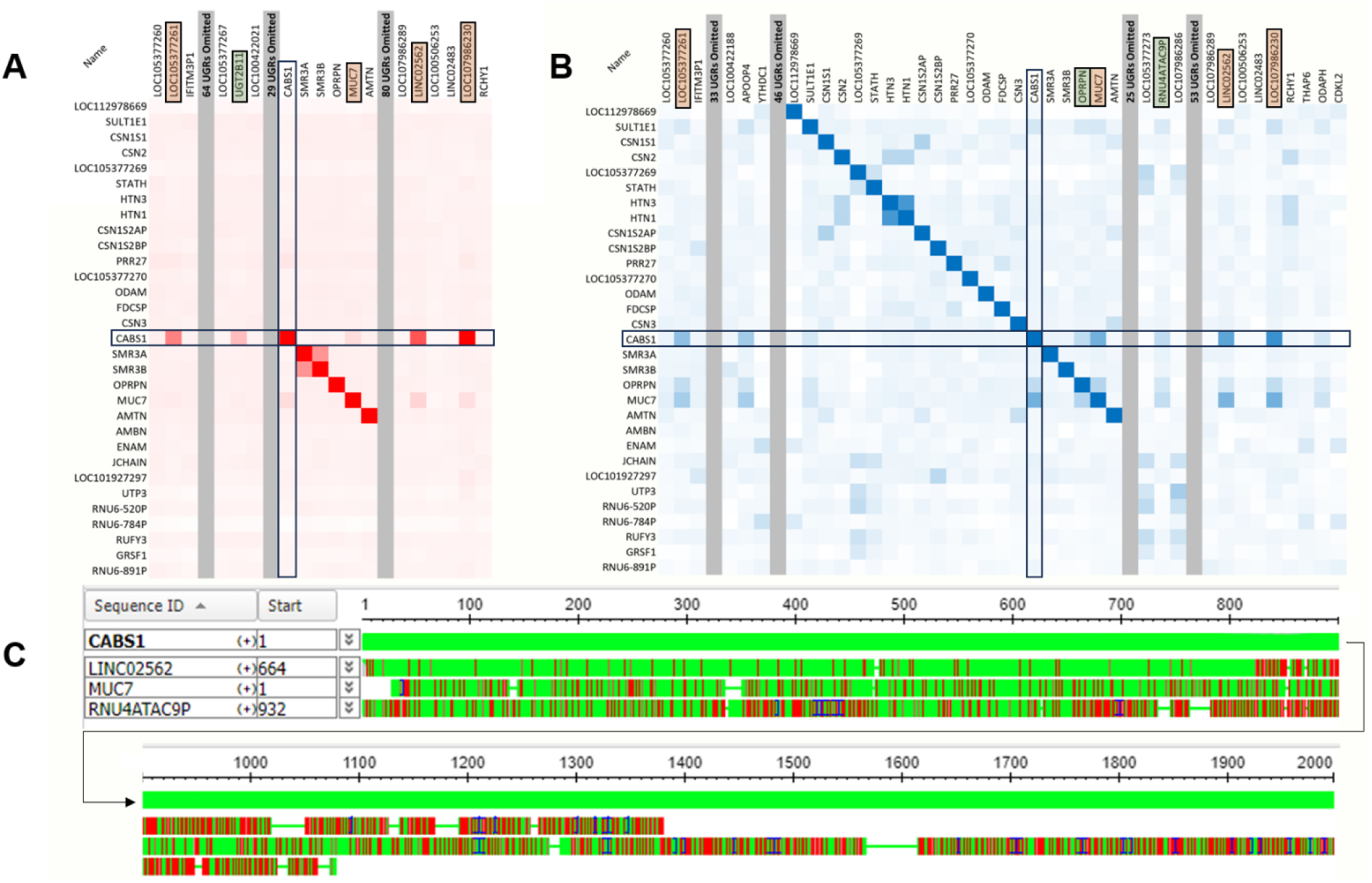
Analysis of the CABS1 neighborhood in Chromosome 4. (A) Partial heatmap with section breaks generated from our software (red). For the purposes of this figure, we have selectively omitted portions of the linear genome leaving a buffer of 1 UGR before and after a similar UGR to CABS1. The grey bars depict the omissions in the linear genome. The first break between IFITM3P1 and LOC105377267 is from nucleotide 66094526 to 68300322. The other two breaks are [69216529 to 70334981] and [70532743 to 75037055]. (B) The same region was validated with Clustal Omega. The omissions for Panel B are as follows: [66094526 to 66003201], [68350090 to 68300322], [70532743 to 72953850], and [72979588 to 75037055]. The CABS1 UGR showed high similarity to a few adjacent UGRs for MUC7, RNU4ATAC9P, and LINC02562, as depicted by the more intense red or blue colour. CABS1 UGR also showed similarity to more distal UGRs of undefined genes (LOC genes). New gene UGRs with relatively high similarity were discovered upon validation with the less rigorous Clustal Omega algorithm such as OPRPN and RNU4ATAC9P and conversely, some unique similarities were highlighted using SEQSIM, such as UGT2B11. (C) Clustal Omega multiple sequence alignment showing the islands of homology of the known genes, MUC7, RNU4ATAC9P and LINC02562, in comparison to the CABS1 UGR sequence. Green indicates exact homology and red indicates nucleotide differences.

### 3.3 Interesting Patterns in Other Chromosomes

While our primary focus was on the case study gene CABS1, we extended the comprehensive analysis to generate similarity heatmaps for every chromosome (beyond the scope of this manuscript). The heatmaps for each chromosome revealed intriguing patterns of similarity between UGRs in both proximal and distal genes. Of particular interest were gene UGRs that were adjacent to each other, which could indicate potential structural and functional synergies.

Figure 5A, shows a small section of the first 100 UGRs on chromosome 1, with similarity scores shown as a heatmap. Striking patterns of similarity in the first 100 UGRs resembled a series of diagonal slashes (arrows in Fig 5B). Additionally, we observed more intriguing patterns such as the checkerboard-like background (open arrow heads point at both the light and dark segments of the checkerboard) emphasized in Figure 5B. These patterns, though distinct from the diagonal slashes, may inform latent genomic interactions. In Figure 5C, distinct bright red (i.e.: very similar UGRs) clusters of similarity among adjacent gene UGRs were also observed. One example is shown with the closed arrowhead in 5B. It’s interesting to note that when highly similar adjacent UGRs form clusters, other UGRs around the islands are dissimilar. This is reminiscent of how topologically associated domains (TADs) form physical loops in chromatin and interact at a higher frequency within the TAD versus adjacent TADs. These results may illustrate intricate genomic interactions to be further elucidated with future research.

**Figure 5.**
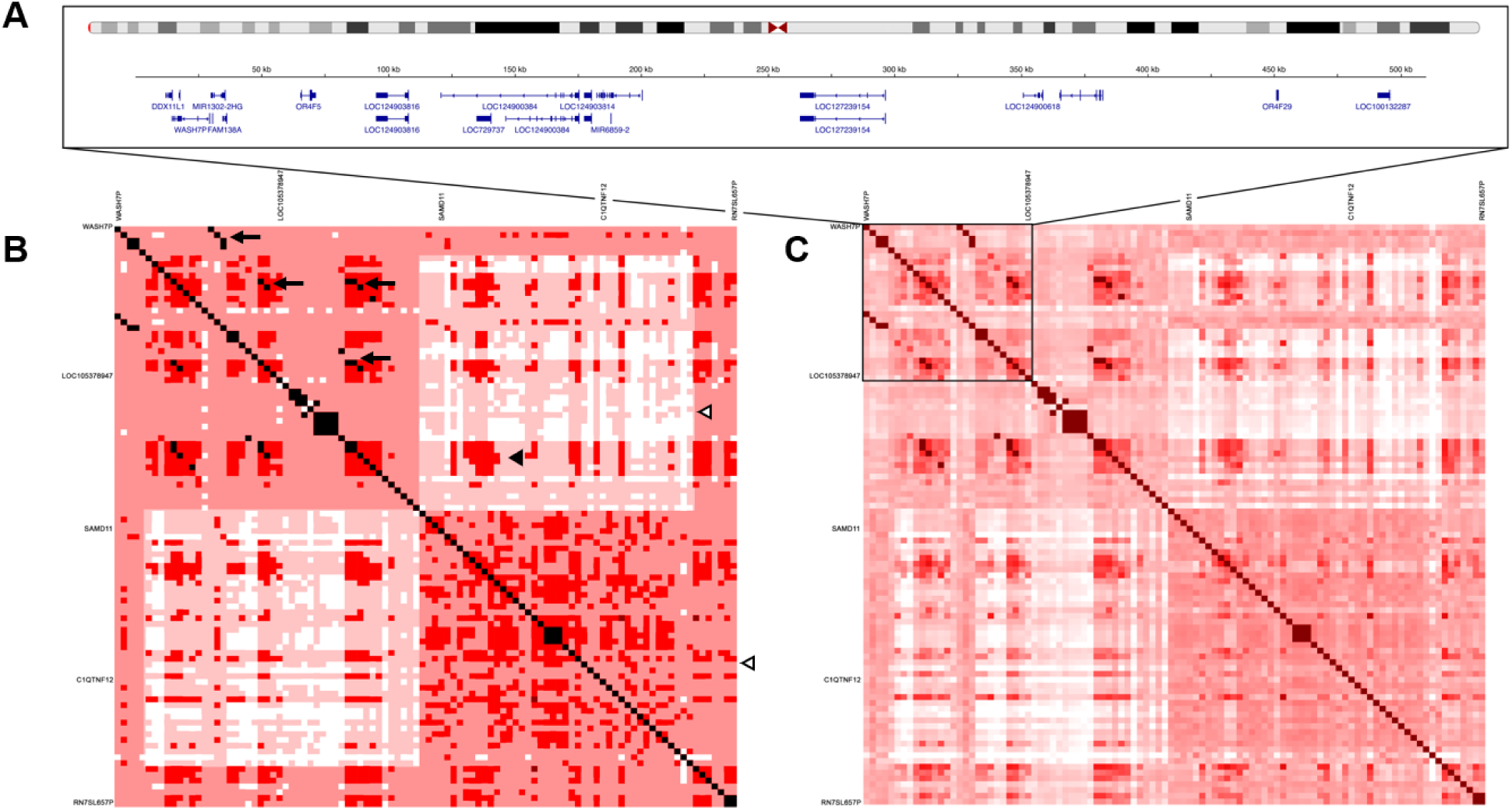
Heatmap data visualization for the first 100 gene UGRs of Chromosome 1. (A) Gene view of the first 500 kb of Chromosome 1 and a segment included in our heatmaps. In this segment, there are 27 genes. (B) Overemphasized graphical representation of the patterns that can be extracted from the heatmap depicted in 5C. The black arrowhead indicates one cluster of homology that is of interest due to the overall pattern created in the map. The white arrowheads indicate a darker and lighter red area observed in the first 100 genes. The black arrows show the diagonal slashes observed in the heatmap. The darker red indicates higher similarity in the UGRs while the light red area depicts less similarity to adjacent UGRs. White areas indicate little to no similarity. (C) The actual heatmap without digital enhancement generated using our novel comparison software and algorithm.

## 4. Discussion

Our research on UGRs has uncovered intriguing patterns of sequence similarity that suggest potential regulatory mechanisms and 3D chromatin conformations. SEQSIM generated a large matrix of pairwise similarity scores between UGRs of every human gene. We discovered 41 novel clusters of UGRs that can be categorized according to features such as transcription factor binding site motifs and segments of transposable elements within them. Our findings align with and expand on existing literature(3,22–24) that will help elucidate mechanisms underlying chromosomal architecture and gene regulation.

Recent research has demonstrated the importance of UGRs, highlighting their involvement in critical conditions like cancer, diabetes and neurodegenerative disorders(25). These UGRs, when mutated or varied, can alter gene expression significantly. Additionally, the role of epigenetic modifications in changing the accessibility of these UGRs to regulatory proteins is a crucial factor in gene expression(26,27). Most research focuses specifically on genes and not the UGRs, highlighting a substantial gap in understanding how UGRs behave under different epigenetic states and how this might affect gene expression, regulation, and even the 3D genome. Specifically, SEQSIM revealed patterns of UGR similarity with adjacent genes that may infer 3D chromatin conformation. SEQSIM also identified previously undetected patterns within the chromosomes, surpassing the data available in existing 3D genome databases(28). The role of UGRs in chromosomal architecture, such as TADs or chromosomal territories that facilitate specific gene interactions, remains poorly understood. It is possible that UGRs play a role in establishing or maintaining these domains, but research in this area is limited. Our research advances the field by providing a tool for genome-wide analysis and comparison of UGRs, and some initial examples of the types of interactions that may occur.

The examination of promoter sequence similarity is essential in elucidating gene regulatory mechanisms and expression patterns. Rhoads and McIntosh showed that genes like the salicylic acid-inducible alternative oxidase gene *aox1* and those involved in disease resistance share similar sequences in their promoters(29). This suggests they might be controlled in similar ways or have evolved together, which is important for determining how genes respond to environmental changes or stress. Likewise, another study focused on genes in *Brassica napus* pollen. The researchers found that these genes have conserved sequences in their promoter regions, which play roles in when and where these genes are active(30). Utilizing SEQSIM, we discerned clusters of similar UGRs in the human genome, which helps identifying potentially co-regulated genes and their associated regulatory elements within each cluster.

Statistical analysis of modularity revealed 41 clusters of UGRs based on sequence similarity, with 67.4% of linked UGRs appearing in the main 5 clusters (Figure 3B). This could indicate 5 main “classes” of UGRs, with several smaller sub-types representing the other 36 clusters (Figure 3C). As mentioned previously, Gagniuc et al. attempted to categorize promoters into 10 generic classes based on sequence features and associated regulatory proteins, and our findings suggest there are indeed different categories or classes of UGRs(3). However, no other researchers, to our knowledge, have attempted to categorize UGRs and promoters based on sequence similarity genome wide. Unique sets of genes associated with different biological pathways, functions, or cellular processes may be represented by these distinct clusters. For example, the identification of protein-coding genes with the highest SEQSIM score to the *CABS1* UGR and within the *CABS1* cluster, such as *VWCE, SPOCK1, THSD4, RNF39* and *TMX2*, suggests shared regulatory mechanisms and potential strategies to interrogate the regulation and function of this poorly known, but apparently widely distributed gene in human tissues(13,31). Extrapolating from other members of its clusters may help us define the functional role of CABS1 and understand its roles in various biologically processes. *SPOCK1* transcripts, for example, are also widely expressed in the human body but the protein itself is expressed in testis, much like CABS1(32). SEQSIM analysis placed the CABS1 UGR in the second largest cluster. The biological functions of the cluster’s 385 members were heterogenous leading to little concrete insight; however, our analysis may still indicate that certain genes with highly similar UGRs work together even if they are not located on the same chromosome. In the future, studies could be conducted to determine the likelihood that genes within the same UGR cluster are co-expressed, however that study is out of scope for this paper. Experts agree that genes positioned closely on the same chromosome tend to co-express or co-regulate more frequently than genes on different chromosomes. However, our findings imply that factors beyond physical proximity, such as sequence similarity and shared control signals that extend across the genome, could influence gene coregulation and co-expression. For example, these long-range interactions could be a complex network of transcription factors that interact specifically to the similar sequences in each of the aforementioned genes.

In our research, we also explored similarities between UGRs on the same chromosome as CABS1, and we also found evidence of functional sites. In Figure 4B, the multiple sequence alignment of the *CABS1* UGR with adjacent UGRs with high similarity, *LINC02562, MUC7* and *RNU4ATAC9P* UGRs, reveals small sections of identical sequences, which could correspond to transcription factor binding site motifs (5 to 20 nucleotides long), promoters, enhancers, or other regulatory elements. CABS1 is primarily expressed in testicular tissues and is thought to be involved in the late stages of spermatogenesis, and it also has been found in human saliva(13). Interestingly, MUC7 (mucin-7, secreted) is a member of the mucin family of proteins which is specifically known for its role in saliva, contributing to the innate defense in the oral cavity. *LINC02562* (long intergenic non-protein coding RNA 2562) is part of a class of RNAs that, while not coding for proteins, can also affect gene regulation. Long non-coding RNAs are known to be involved in a variety of biological processes, including chromatin remodeling, transcriptional control and post-transcriptional modifications(33,34). Similarly, the small nuclear RNA U4atac Pseudogene 9 (*RNU4ATAC9P*) may be involved in minor spliceosome function. Traditionally thought to be non-functional, recent studies have suggested that pseudogenes, and types of pseudogenes like retrotransposons, may play a regulatory role in gene expression(34,35). The similarity of the CABS1 UGR to these other UGRs could provide clues to how CABS1 is regulated in the body and perhaps some of its functions.

The history of CABS1 research is intertwined with studies on the rat gene *Vcsa1* (absent in humans) and its protein, submandibular rat 1 (SMR1), which have significant roles in inflammation, shock response, and neuroprotection(12). Specifically, human CABS1 shares a similar amino acid sequence near the carboxyl terminal (TDIFELL) with rat SMR1 (TDIFEGG), and both sequences have anti-inflammatory activities(12). On chromosome 4, *CABS1* is surrounded by other genes with activities similar to rat *Vcsa1*, so it was interesting that the UGR of *CABS1* did not have high similarity to that of related adjacent genes, *SMR3A* or *SMR3B* that like CABS1 are associated with male reproductive function(36–38). Different UGRs in genes with potentially similar biological activities could be a result of evolutionary divergence and the need for tissue-specific expression. This diversity may help the same or similar functions to be carried out in different tissues, developmental stages, or environmental conditions.

Some UGRs in the CABS1 group contain parts of a long interspersed nuclear element (LINE), a type of transposable element (TE)(39,40). TEs are DNA sequences capable of moving or “jumping” within the genome and can insert themselves into new genomic locations where they may act as regulatory elements by providing binding sites for transcription factors or even altering chromatin structure(41,42). This alteration of chromatin structure by TEs plays a role in shaping the 3D architecture of the genome. TEs can lead to the formation of new chromatin loops and domains, that affect the proximity of regulatory elements to target genes, impacting transcription and genomic structure(43).

Furthermore, distinct TE subfamilies that function as tissue-specific enhancers in colon and liver cancers have been identified(44). These TEs are characterized by genomic features associated with active enhancers, such as epigenetic marks and transcription factor binding, and are associated with differentially expressed genes in these cancer types. The presence of the large islands of identical sequences (green regions, Fig 4C) in the multiple sequence alignments suggests that these features are evolutionarily conserved in the UGR and may hold functional importance. Variations or mutations in these conserved regions may cause gene dysregulation and abnormal phenotypes. A potential future application of SEQSIM could be the comparative analysis of UGRs among individuals or populations to identify genetic variants associated with increased susceptibility to dysregulation. Integrative analysis of 111 reference human epigenomes has shown the relationship between histone modification patterns, DNA accessibility, DNA methylation, and RNA expression(45). This wealth of epigenomic data can be leveraged to interpret the molecular basis of human disease and further understand the regulatory mechanisms and functional importance of UGRs. It would also be interesting to study whether the TEs responsible for these large islands of homology are conserved across species, and to determine whether they retained their historical regulatory functions or acquired new roles over evolution(46,47).

Recent studies have begun to shed light on the intricate relationship between 3D chromatin architecture and gene expression under various conditions. For instance, research on primate cortical neurons exposed to morphine showed rearrangement in chromosome territories and alterations in TADs, which were associated with changes in gene expression(48). In the pursuit of deciphering 3D chromatin architecture, methodologies such as chromosome conformation capture (3C) and its derivative Hi-C, as well as techniques like DNA-FISH, have been instrumental. These techniques, while revolutionary, have presented researchers with other issues: the need to navigate through vast amounts of data generated and the need to analyze this data to elucidate chromosomal interactions and their implications in gene regulation with precision and accuracy. The development of new software tools, like SEQSIM, to facilitate this analysis is an important step towards achieving these goals(49,50).

As mentioned previously, the observed patterns of similarity among adjacent gene UGRs in our heatmaps enrich current 3D genome databases by uncovering data in previously undocumented loci. While the exact functional significance of these patterns remains to be elucidated, several interpretations can be proposed. Firstly, the patterns of similarity observed in the heatmaps might indicate the existence of distinct chromosomal neighborhoods or domains within a single chromosome. Genes within the same neighborhood may experience similar chromatin environments, accessibility to regulatory elements, or proximity to nuclear compartments involved in gene expression. A recent study demonstrated that 3D chromatin architecture in T-cells facilitates cell-specific factor binding and works synergistically with global and tissue-specific regulators, such as CTCF (CCCTC-binding factor) and SATB1 (special AT-rich Sequence Binding Protein 1), to control gene expression(51). The identified UGR patterns could indicate discrete functional regions within a chromosome, where neighboring genes work in concert to fulfill specific cellular functions, akin to conventional TADs. Similar adjacent gene co-regulation, where functionally related genes are spatially clustered in the chromosome, exists in yeast. Genes involved in ribosome biogenesis, for example are found to be clustered, suggesting their spatial arrangement facilitates coordinated expression(52,53). Adjacent genes within the same TAD may share similar chromatin states, including histone modifications, DNA methylation patterns, or nucleosome occupancy(54). Intriguingly, when juxtaposing our findings with conventional Hi-C 3D genome data, we observed patterns unique to our SEQSIM heatmaps. Although Hi-C has been instrumental in revealing the 3D organization of the genome, it may be limited by resolution and may not capture all potential regulatory interactions, especially those that are transient or condition specific. The unique relationships shown in SEQSIM may help fill these gaps in Hi-C data, and exploring these relationships may uncover novel gene networks or pathways involved in specific biological processes or diseases.

Lastly, the identified patterns could also possess evolutionary significance. Adjacent UGRs with conserved similarity patterns across species may indicate functional constraints or co-evolutionary relationships. This is supported by research showing that certain areas near genes, known as cis-regulatory regions, are under evolutionary pressure to either stay the same or change in particular ways, especially in humans(55). Moreover, comprehensive studies have identified 101 specific regions within 100 kb of known genes in the human genome that have recently undergone adaptive evolution, including genes in the immune system and heat shock genes(56). These regions are subject to both stabilizing and directional selection, indicating their critical role in gene regulation, and maintaining genomic stability or facilitating adaptive processes. Further evolutionary studies are warranted to elucidate the conservation of these patterns and investigate their role in maintaining genomic stability or facilitating adaptive processes.

While our speculations provide a foundation for hypothesis generation, they require more in-depth computational analysis and experimental validation. Currently, we do not have a method to conduct a full cluster analysis on the entire genome. However, we plan to explore artificial intelligence techniques and other iterative computational approaches to enable the processing of big data. Additionally, further research utilizing complementary techniques, such as functional genomics, chromatin conformation capture assays, and gene perturbation studies, can shed light on the functional implications of these observed patterns and provide a deeper understanding of the intricate organization and regulation of genes within chromosomes.

## 5. Conclusion

In conclusion, our research demonstrates the utility of bioinformatics tools, specifically SEQSIM, for the comprehensive analysis and comparison of UGRs across the human genome. Our novel approach revealed distinct UGR clusters, indicating potential co-regulation, unique gene sets associated with different biological pathways, and the potential presence of evolutionary conserved regions. Notably, identical sequence segments may be critical regulatory elements or TE, implying an intricate interplay of genomic elements in gene regulation.

While computational constraints limited the depth of our analysis, the observed similarity patterns among adjacent gene UGRs revealed in our heatmaps have sparked new hypotheses about co-regulation, chromosomal organization, and evolutionary significance. Future experimental validation and the use of additional techniques will shed light on these observations, elucidating the complexities of genome organization and regulation thereby enriching our understanding of human health and disease.

## Acknowledgements

We thank Gilbert Lee for providing technical support for this project.

## Funding Information

This work was supported by the Natural Sciences and Engineering Research Council of Canada [RGPIN-2020-04553].

## Notes

### Competing Interest Statement

The authors have declared no competing interest.

